# iterb-PPse: Identification of transcriptional terminators in bacterial by incorporating nucleotide properties into PseKNC

**DOI:** 10.1101/2020.01.17.910232

**Authors:** Yongxian Fan, Wanru Wang, Qingqi Zhu

## Abstract

Terminator is a DNA sequence that give the RNA polymerase the transcriptional termination signal. Identifying terminators correctly can optimize the genome annotation, more importantly, it has considerable application value in disease diagnosis and therapies. However, accurate prediction methods are deficient and in urgent need. Therefore, we proposed a prediction method “iterb-PPse” for terminators by incorporating 47 nucleotide properties into PseKNC- `I` and PseKNC- `II` and utilizing Extreme Gradient Boosting to predict terminators based on *Escherichia coli* and *Bacillus subtilis*. Combing with the preceding methods, we employed three new feature extraction methods K-pwm, Base-content, Nucleotidepro to formulate raw samples. The two-step method was applied to select features. When identifying terminators based on optimized features, we compared five single models as well as 16 ensemble models. As a result, the accuracy of our method on benchmark dataset achieved 99.88%, higher than the existing state-of-the-art predictor iTerm-PseKNC in 100 times five-fold cross-validation test. It’s prediction accuracy for two independent datasets reached 94.24% and 99.45% respectively. For the convenience of users, a software was developed with the same name on the basis of “iterb-PPse”. The open software and source code of “iterb-PPse” are available at https://github.com/Sarahyouzi/iterb-PPse.

## 1 Introduction

DNA transcription is an important step in the inheritance of genetic information and terminators control the termination of transcription which exists in sequences that have been transcribed. When transcription, the terminator will give the RNA polymerase the transcriptional termination signal. Identifying terminators accurately can optimize the genome annotation, more importantly, it has great application value in disease diagnosis and therapies, so it is crucial to identify terminators. Whereas, using traditional biological experiments to identify terminators is extremely time consuming and labor intensive. Therefore, a more effective and convenient began to be applied in researches, that is, adopting machine learning to identify gene sequences.

Previous research found there are two types of terminators in prokaryotes, namely Rho-dependent and Rho-independent[1], as shown in Fig 1. Although there have been a lot of studies on the prediction of terminators, most of them only focused on one kind of them. In 2004, Wan XF, Xu D et al. proposed a prediction method for Rho-independent terminators with an accuracy of 92.25%. In 2005, Michiel J. L. de Hoon et al. studied the sequence of Rho-independent terminators in *B. subtilis*[2], and the final prediction accuracy was 94%. In 2011, Magali Naville et al. conducted a research on Rho-dependent transcriptional terminators[3]. They used two published algorithms, Erpin and RNA motif, to predict terminators. The specificity and sensitivity of the final results were 95.3% and 87.8%, respectively. In 2019, Macro Di Simore et al. utilized the secondary structure of the sequence as a feature[4], the classification accuracy of the Rho-independent terminators was 67.5%. Not like the above experiments Lin Hao et al. studied the prediction of two kinds of terminators in bacterial[5],they developed a prediction tool for terminators with an accuracy of 95% in 2018.

**Fig 1.**
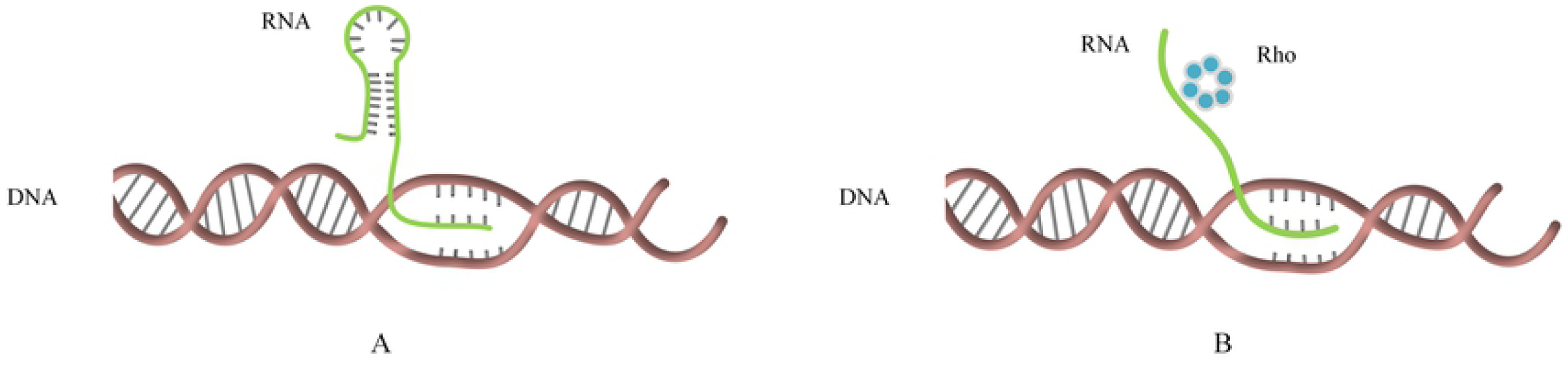
Transcriptional termination process. (A) The termination do not require Rho. The transcription stops when the RNA forms the stem loop structure. (B) The termination dependent on Rho.

To further improve the prediction accuracy, we obtained 503 terminator sequences, 719 non-terminator sequences of *Escherichia coli* (*E. coli*), and 425 terminator sequences, 122 non-terminator sequence of *Bacillus subtilis* (*B. subtilis*) to construct the benchmark dataset and two independent sets. Furthermore, we proposed three new feature extraction methods (K-pwm, Base-content, Nucleotidepro) to combine them with PseKNC - `I`[6] and PseKNC - `II`[5], then applied the two-step method to select effective features. In addition, we compared five single models (Support Vector Machine (SVM), Naive Bayes, Logistic Regression (LR), Decision Tree, Multi-layer Perceptron (MLP), K-Nearest Neighbor (KNN)) as well as 16 ensemble models based on AdaBoost, Bagging, Extreme Gradient Boosting (XGBoost) and Gradient Boosting Method (GBM). Finally, we proposed a prediction method “iterb-PPse” for terminators.

## 2 Materials and Methods

As shown in the Fig 2, our study is mainly divided into the following steps[7]: (1) data collection, (2) feature extraction, (3) feature combination, (4) feature selection, (5) classification, (6) result evaluation, (7) prediction method.

**Fig 2.**
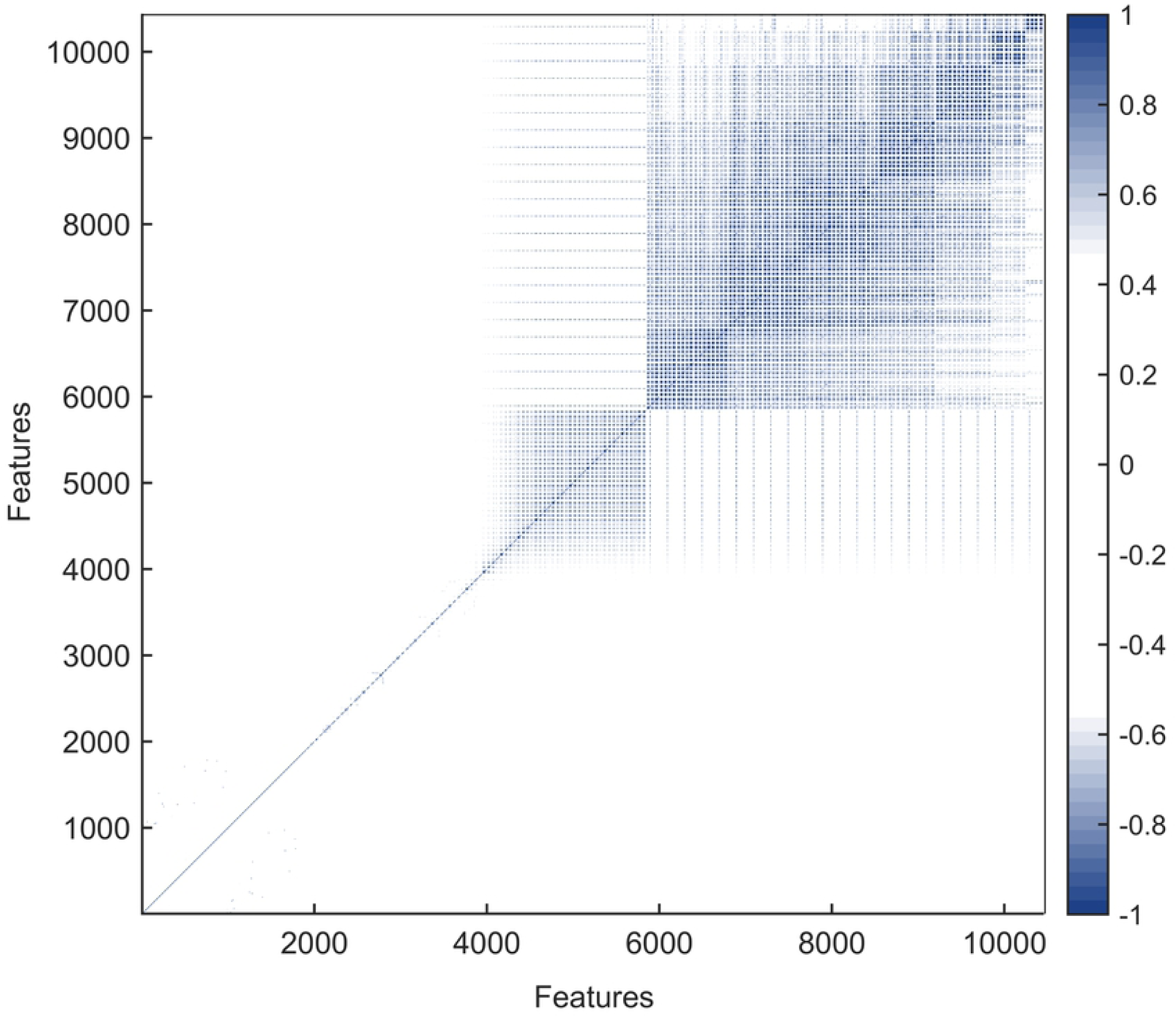
The overall framework. A shows main steps of our study. First step is using five extraction methods to deal datasets, then select more important features by two-step feature selection method, finally compared different models using the selected features. The “iterb-PPse” is the method we proposed to predict terminators. B illustrates the prediction process of “iterm-PPse”. It extracts three features from gene sequences at first, namely Pse5NC-`I`, Pse5NC-`II`, 47 nucleotide properties. Then sort all features using F-score and select the best feature set by IFS. Finally utilizes trained XGBoost to determine whether these sequences are terminators.

### 2.1 Data Collection

In our study, the initial datasets were obtained from http://lin-group.cn/server/iTerm-PseKNC [2], which includes 280 terminator sequences, 560 non-terminator sequences of E. coli, and 425 terminator sequences of B. subtilis. To generate reliable benchmark dataset and independent dataset, we collected another 76 terminator sequences, 159 non-terminator sequences from *E. coli K-12* genome in the database RegulonDB[8], and 122 non-terminator sequences of *B. subtilis* were gathered from database DBTBS[2, 9]. The non-terminator sequences of *E. coli* were intercepted from −100 bp to −20 bp upstream and 20 bp to 100 bp of positive samples not used in the benchmark dataset. The non-terminator sequences of *B. subtilis* were intercepted from −102 bp to −20 bp upstream and 20 bp to 102 bp of positive samples. At last, we divided the collected sequences into the benchmark set and the independent dataset at a ratio of 8: 2. In order to accurately evaluate the identification accuracy of our method to different bacteria, we divided the independent test set into two. Details of the benchmark dataset and independent sets are shown in Tables 1 and 2 of respectively. All sequences of *E. coli* and *B. subtilis* could be found in S1-S7 Tables of Supplementary data.

**Table 1.** Benchmark dataset.

| Species | Category | Number | Length |
| --- | --- | --- | --- |
| *E. coli* | Rho-dependent terminator | 18 | ~50 bp |
|  | Rho-independent terminator | 385 | ~50 bp |
|  | non-terminator | 575 | 80 bp |
| *B. subtilis* | Rho-independent terminator | 340 | ~50 bp |
|  | non-terminator | 98 | 82 bp |

**Table 2.** Independent dataset.

| Species | Category | Number | Length |
| --- | --- | --- | --- |
| *E. coli* | Rho-independent terminator | 100 | ~50 bp |
|  | non-terminator | 143 | 80 bp |
| *B. subtilis* | Rho-independent terminator | 85 | ~50 bp |
|  | non-terminator | 24 | 82 bp |

### 2.2 Feature extraction

How to extract effective features from DNA sequences is a particularly important step. At present, the input of most machine learning methods must be numerical values rather than character sequences[10], such as decision tree, logistic regression etc. Thus, it is essential to make use of proper feature extraction methods to represent sequences.

#### 2.2.1 K-pwm

The new feature extraction method “K-pwm” mainly employed the Position Weight Matrix[11-14], where K represents *k*-tuple nucleotides. Considering that the length of negative samples is different from that of the positive samples in the benchmark set. we made a little modification to the calculation of the final sequence score to eliminate the negative impact of sequence length. A total of 6 feature sets were obtained by using this method, namely the position weight features corresponding to *k* =1, 2, 3, 4, 5, 6. The calculation steps are shown below.

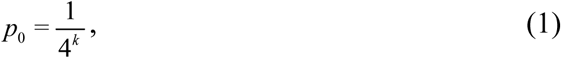

where *p*_0_ represents the background probability of the occurrence of *k*-tuple nucleotides.

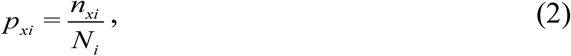

where *p*_*xi*_ indicates the probability of *k*-tuple nucleotide *x* appearing at site *i*.

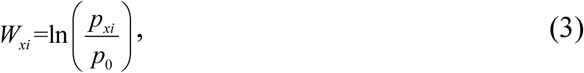

where *W*_*xi*_ is the element in the position weight matrix.

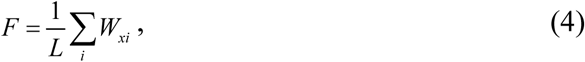

where *L* is the length of the corresponding sequence.

#### 2.2.2 Base-content

Given that the rho-independent terminators are rich in GC base pairs, we extracted a set of features and collectively referred to as Base-content[15, 16]. Specifically, we mainly obtained the content features of the single nucleotide(A, C, G, T) in each DNA sequence[17, 18]. In this paper, 5 kinds of base content features(atContent, gcContent, gcSkew, atSkew, atgcRatio)[15, 16, 19-21] were took into account.

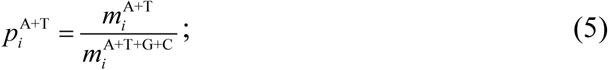

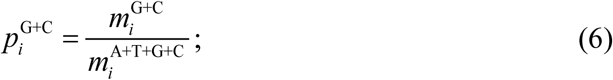

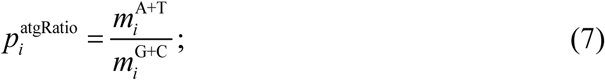

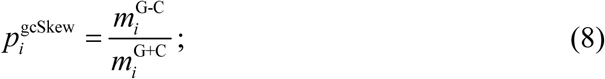

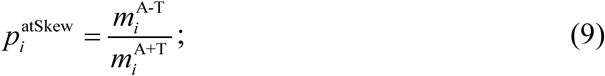

where *m*G *i, m*C *i*are the contents of G and C in the *i*-th sequence, respectively. *m*A+T *i, m*G+C *i, m*A+T+G+C *i* are the contents of “A+T”, “G+C” and “A+T+G+C”, respectively. *m*A-T *i, m*G-C *i* represent the content of “A-T” and “G-C”, respectively.

#### 2.2.3 Nucleotidepro

Nucleotide properties of DNA sequences play a key role in gene regulation[22]. Therefore, we proposed a new feature extraction method “Nucleotidepro” involving 47 properties[23] not covered previously, including 3 nucleotide chemical properties[24], 32 dinucleotide physicochemical properties and 12 trinucleotide physicochemical properties.

To extract corresponding features, we employed a 47\**L* dimension matrix to represent each sequence. *L* is the length of the corresponding sequence. As shown in the Table 3, we used 0 and 1 to represent the chemical properties of different nucleotides. Then we iterated through each sequence and assigned the values of different properties for different nucleotide to the corresponding elements in the matrix. The nucleotide properties and corresponding standard-converted values[23] for the 47 properties can be obtained from the Tables S8 and S9 from Supplementary data.

**Table 3.** Corresponding values for different chemical properties.

| Chemical | Category | Nucleotides | Value |
| --- | --- | --- | --- |
| Ring structure | Purine | AG | 0 |
|  | Pyrimidine | CT | 1 |
| Hydrogen bond | Strong | CG | 0 |
|  | Weak | AT | 1 |
| Functional group | Amino | AC | 0 |
|  | Keto | GT | 1 |

#### 2.2.4 PseKNC-I

PseKNC-I [6] is generally understood to mean the parallel correlation PseKNC. It combines K-tuple nucleotides components [25] with 6 physicochemical properties [22] (rise, slide, shift, twist, roll, tilt), not only considering the global or long-range sequence information, but also calculating the biochemical information of DNA sequences. The PseKNC-`I` features can be obtained directly through the online tool Pse-in-one [26, 27], or run our code to process multiple sequences at the same time.

By changing the value of *K*, more features could be obtained. However, as the dimension of the feature matrix increases, it may lead to over-fitting and generate a large amount of redundant data[28]. Therefore, only three feature sets were extracted when *K* = 4, 5 and 6, respectively.

#### 2.2.5 PseKNC-II

PseKNC-`II`, also known as the series correlation PseKNC[5]. PseKNC-`II` also calculated the K-tuple pseudo nucleotide properties, but unlike PseKNC-I, it considered the difference between properties. By changing the value of *K*. We extracted three feature sets when *K*= 4, 5, 6 respectively.

### 2.3 Feature combination

Each feature extraction method can extract distinctive features of the DNA sequence with different emphasis. To further optimize the prediction results, we analyzed the performance of five feature extraction methods by training XGBoost to predict terminators and selected the more effective features from each method to combine. The specific combination method will be introduced in the section **Results**.

### 2.4 Feature selection

Feature selection is an important data process, which could not only reduce the computation time, but also remove redundant data, and select more effective features, finally greatly improve the prediction accuracy[28].Hence, the two-step method was adopted to select features.

#### 2.4.1 Feature analysis

To present the correlation between features, the Pearson correlation coefficients were calculated to construct correlation matrix. If the two properties change in the opposite direction, it is a opposite effect. As shown in Fig 3, the features contain some redundant data, so it is necessary to utilize the two-step feature selection method[5, 17, 29].

**Fig 3.**
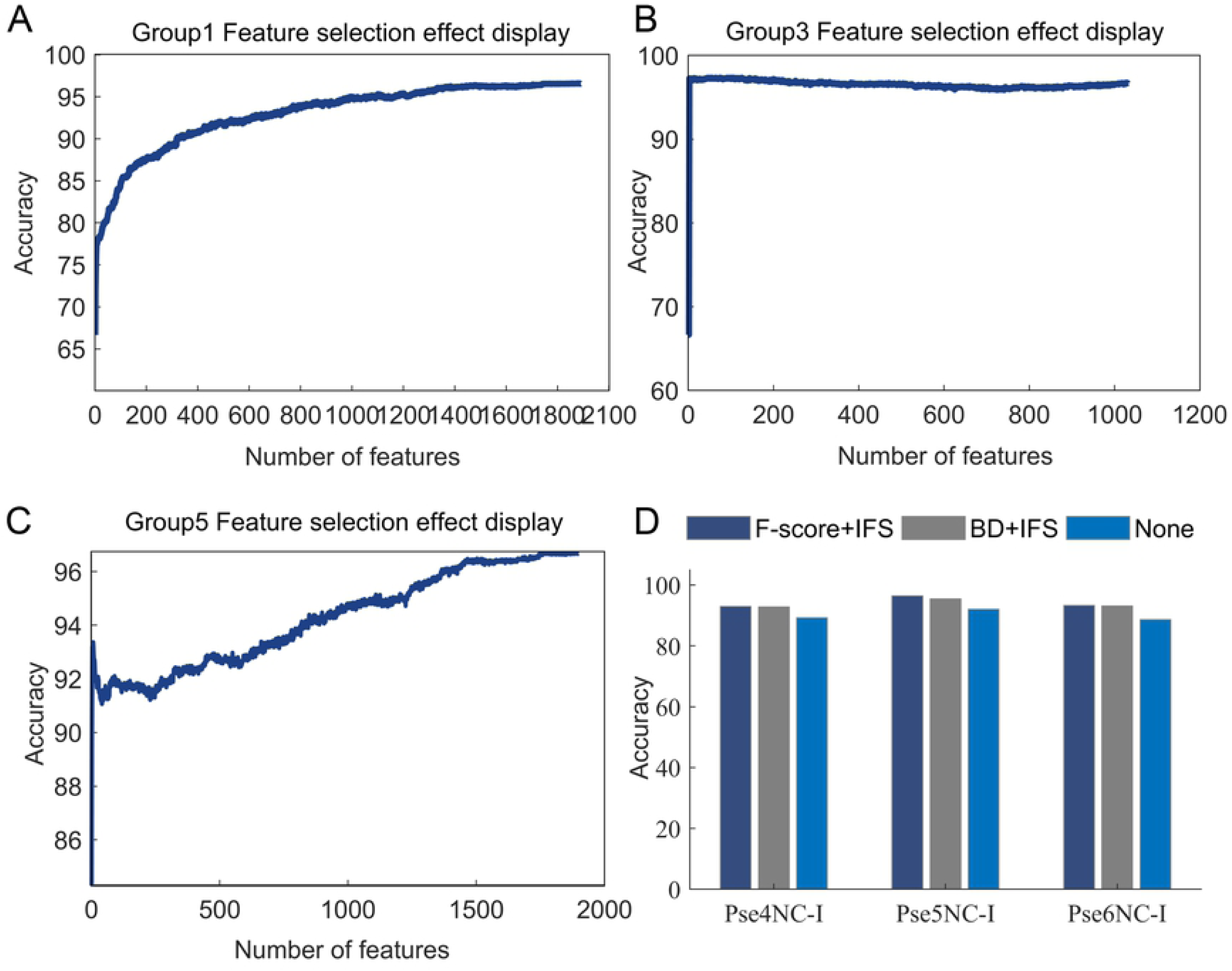
Correlation of all features. The correlation between all features obtained by calculating the Pearson correlation coefficient.

#### 2.4.2 Feature Sorting

The first step is utilizing feature sorting methods. The main task of feature sort is to analyze the importance of each feature for prediction of terminators. The top features are more helpful in predicting terminators.

##### F-score

F-score[6] is a method for measuring the ability of a feature to distinguish between two classes. Given the training set *x*, if *n*^+^ and *n*^-^ stand for the number of positive and negative samples, respectively. The F-score of the *i*-th feature is inferred to be:

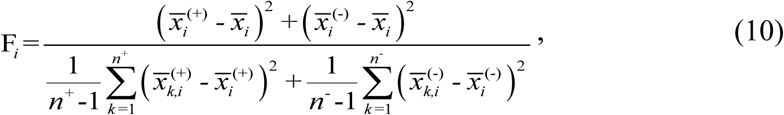

where 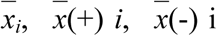 represent the average of the *i*-th feature in all samples, positive samples, and negative samples, respectively. 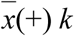, *i* is the *i*-th feature of the *k*-th positive sample, 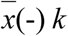,*i* is the *i* -th, feature of the *k*-th negative sample. The larger the F-score, the more distinctive this feature. The existing feature sorting toolkit fselect.py can be obtained from http://www.csie.ntu.edu.tw/~cjlin/.

##### Binomial distribution

As well as, binomial distribution[27, 30] were used to sort the features[31, 32]. The specific process is as follows:

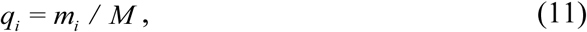

where *q*_i_ is the prior probability, *m*_*i*_ represents the number of *i*-th samples (*i* =1,2 indicates positive and negative respectively), and *M* is the number of all samples.

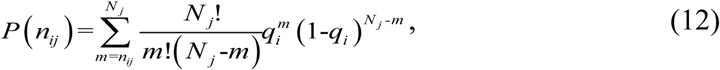

where *n*_*ij*_ represents the times of the *j*-th feature appears in the *i*-th samples, and *N*_*j*_ is the times of the *j*-th feature appears in all samples.

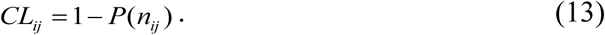

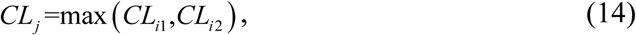

where *CL*_*j*_ is the confidence level, the higher the confidence level, the higher the credibility. Therefore, the confidence level of each feature was ranked in descending order according to the corresponding *CL*_*j*_.

#### 2.4.3 Incremental feature selection

The second step is Incremental Feature Selection(IFS)[33]. It uses a feature as the training set at first, then the sorted features are added to the training set one by one, finally find the number of features corresponding to highest classification accuracy.

### 2.5 Data normalization

It is necessary to process the data into the required format before conducting experiments, such as normalized. Our study first employed function “mapminmax” for data normalization, its purpose is to make data limited in a certain range, such as [0, 1] or [-1, 1], thereby eliminating singular sample data leading to negative impact.

In addition, it should be noted that data normalization is not applicable to all classification algorithms, and sometimes it may lead to a decrease in accuracy. Data normalization applies to optimization problems like AdaBoost, Support Vector Machine, Logistic regression, K-Nearest Neighbor but not probability models such as decision tree.

### 2.6 Model

#### 2.6.1 Single model

##### SVM

The principle of SVM[34] is using a series of kernel functions to map the initial feature sets to high-dimensional space, and then finding a hyperplane in high-dimensional space to classify samples. The SVM pattern classification and regression package LIBSVM is available at https://www.csie.ntu.edu.tw/~cjlin/libsvm/oldfiles/.

##### Naïve Bayes

Naïve Bayes uses the prior probability of an object to calculate posterior probability belongs to one of the categories by using the Bayes formula. The object belongs to the class whose corresponding posterior probability is the greatest.

##### LR

LR usually utilizes known independent variables to fit the model *y*=*w*^T^*x*+*b*. Then, predict the value of a discrete dependent variable (whether true or false). Besides its output value should be 0∼1, so it is very suitable for dealing with the two-class problem.

##### KNN

The main principle of the K-Nearest Neighbor is to find *k* samples closest to the sample to be classified. Then count which category has the largest number of samples, and the current sample belongs to this category.

##### Decision Tree

Decision Tree is based on the tree structure which usually formed by a root node, several leaf nodes and some branches. A node represents an attribute, each branch indicates an option, and each leaf represents a classification result. The principle is to construct a tree with the maximum information gain as a criterion, combine various situations through a tree structure, and then employ it to predict new samples.

##### MLP

MLP with multiple neuron layers, also be known as Deep Neural Networks. Similar to a common neural network, it has an input layer, implicit layers, an output layer, and optimizes the model by information transfer between layers.

#### 2.6.2 Ensemble model

##### Bagging

Bagging’s main principle is to integrate multiple base models of the same kind in order to obtain better learning and generalization performance. Single model SVM, Naïve Bayes, Decision Tree[35] and LR were employed as the base classifier respectively. First, the training set is separated into multiple training subsets to train different models. Then make final decision through the voting method.

##### AdaBoost

AdaBoost is a typical iterative algorithm whose core idea is to train different classifiers (weak classifiers) using the same training set. It adjusts the weight based on whether the sample in each training set is correct and the accuracy of the last round. Then, the modified weights are sent to next layer for training, the classifier obtained by each training are integrated as the ultimate classifier. In our study, Decision Tree, SVM, LR and Naïve Bayes were mainly adopted as the weak classifier for iterative algorithm.

##### GBM

finds the maximum value of a function by exploring it along the gradient direction. The gradient operator always points to the fastest growing direction. Because of the high computational complexity, the improved algorithm only uses one sample point to update the regression coefficient at a time, which greatly improves the computational complexity of the algorithm.

##### XGBoost

XGBoost which utilizes the cart tree that can get the predicted score as the base classifier, optimizes different trees in turn during training, adds them to the integrated classifier, and finally get the predicted scores of all trees. The scores are added together to get the classification results.

### 2.6.3 Parameter Optimization

Before applying various models, we studied the parameters of each model and selected some more important to optimize by grid search using 100 times 5-fold cv scheme[36], as shown in Table 4.

**Table 4.** Parameters and the value range of parameter adjustment.

| Model | Parameter | Value |
| --- | --- | --- |
| SVM | $c, g$ | $[2^{-5}, 2^{15}] \Delta=2, [2^{-15}, 2^{-5}] \Delta=2^{-1}$ |
| LR | $c, solver$ | $[0.1, 1] \Delta=0.1$<br>newton-cg, lbfgs, liblinear, sag |
| MLP | $alpha$ | 0.001, 0.01, 0.1, 0.5, 1, 1.5 |
| Decision Tree | $min\_sample\_split, max\_depth$ | $[2, 30] \Delta=2, [1, 10] \Delta=1$ |
| Bagging | $n\_estimators$ | $[10, 1000] \Delta=50$ |
| AdaBoost | $n\_estimators, learning\_rate$ | $[10, 1000] \Delta=50, [0.1, 1] \Delta=0.1$ |
| GBM | $learning\_rate, n\_estimators$<br>$max\_depth, max\_features,$<br>$random\_state$ | $[0.1, 1] \Delta=0.1, [10, 1000] \Delta=50$<br>$[1, 10] \Delta=1$ |
| XGBoost | $n\_estimators, learning\_rate$ | $[10, 1000] \Delta=50, [0.1, 1] \Delta=0.1$ |

$\Delta$ represents the step size.

### 2.7 Cross-validation test

The 5-fold cross-validation (5-fold CV) can effectively avoid over-fitting and under-learning[37], and the results obtained are more convincing. First randomly divide the dataset into 5 pieces. One of them was employed as the test set and the other four were used as training sets. The above process is repeated until each of the five datasets serves as the test set[38]. Since the datasets are randomly divided, the results are accidental. The stability of the results can be improved by performing repeatedly.

### 2.8 Independent test

To test the prediction performance, we utilized the independent set to test prediction performance of terminators. The initial independent sets were obtained from http://lin-group.cn/server/iTerm-PseKNC [2], containing sequences of *E. coli* and *B. subtilis*, respectively. However, both of them do not include negative samples, which result in the test results are not convincing. Therefore, we collected another 159 non-terminator sequences of *E. coli* and 122 non-terminator sequences of *B. subtilis* from database RegulonDB and DBTBS to construct two reliable independent sets.

### 2.9 Performance measures

For the sake of better presentation and comparison of the experiments results, we mainly calculated the following four evaluation parameters[39-41].

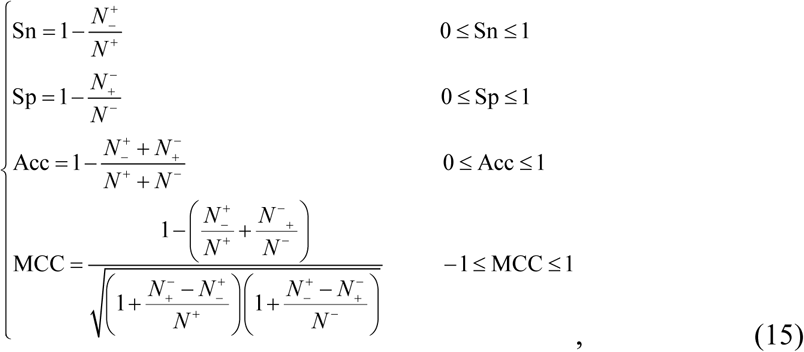

where *N*^+^ represents the number of terminator sequences, and *N*^-^ is the number of non-terminator sequences, *N*+ -indicates the number of positive samples mistaken as negative samples, and *N*-+ indicates the number of negative samples mistaken as positive samples. Sn and Sp delegate the ability of the model to accurately predict samples. Acc reflects the prediction accuracy of models. MCC measures the performance of model[5] on the unbalanced benchmark dataset[42, 43].

In addition to the above four evaluation parameters, the ROC curve was adopted to evaluate the comprehensive performance of different method. It is a comprehensive indicator of continuous variables of sensitivity and specificity. AUC is the area below the ROC curve. Generally, the higher the value of AUC, the higher the classification accuracy[17].

## 3 Results and Discussion

### 3.1 Analysis of feature selection

As shown in Fig 4, we compared the experimental results with and without feature selection, and drew the accuracy corresponding to different number of features after IFS. It is clear that the number of features has a great influence on the classification accuracy, and too many characteristics are bad, so it is necessary to select features. Furthermore, F-score is better than binomial distribution. Therefore, “F-score+IFS” was chose to conduct feature selection.

**Fig 4.**
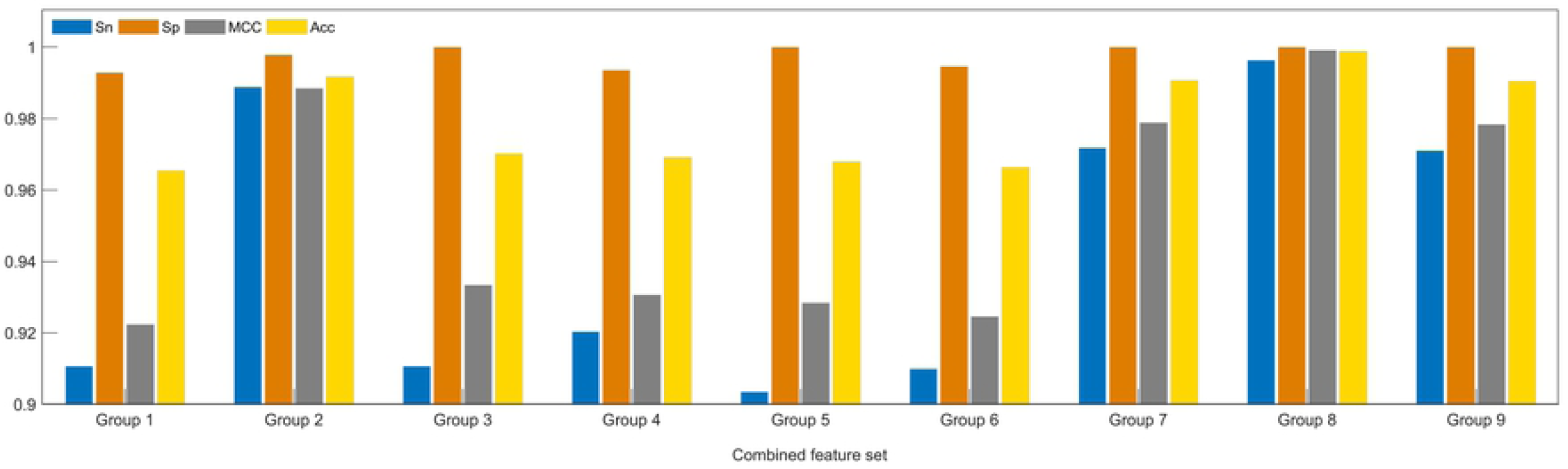
Performance of feature selection. (A)-(C) Relationship between the number of features and classification accuracy of three combined feature sets respectively. (D) Comparison of prediction results using three PseKNC-`I` features and different feature sorting methods. The combined feature set is described in detail in the next section.

### 3.2 Comparison of different feature extraction methods

We compared the performance of different feature extraction methods by training XGBoost to predict terminators. As shown in Fig 5, PseKNC-`I`, PseKNC-`II`, k-pwm, and nucleotidepro are all effective, but the performance of base content is not ideal. Hence, the more effective features were selected to construct combined feature sets. In the end, a total of nine group features were obtained. Details of the combination method are shown in Table 5. As shown in Fig 6, Group 8 stands out in terms of Sn, Sp, MCC and Acc from other combined feature sets. Consequently, the three features Pse5NC-`I`, Pse5NC-`II`, 47 nucleotide properties were applied to formulate all samples.

**Table 5.**
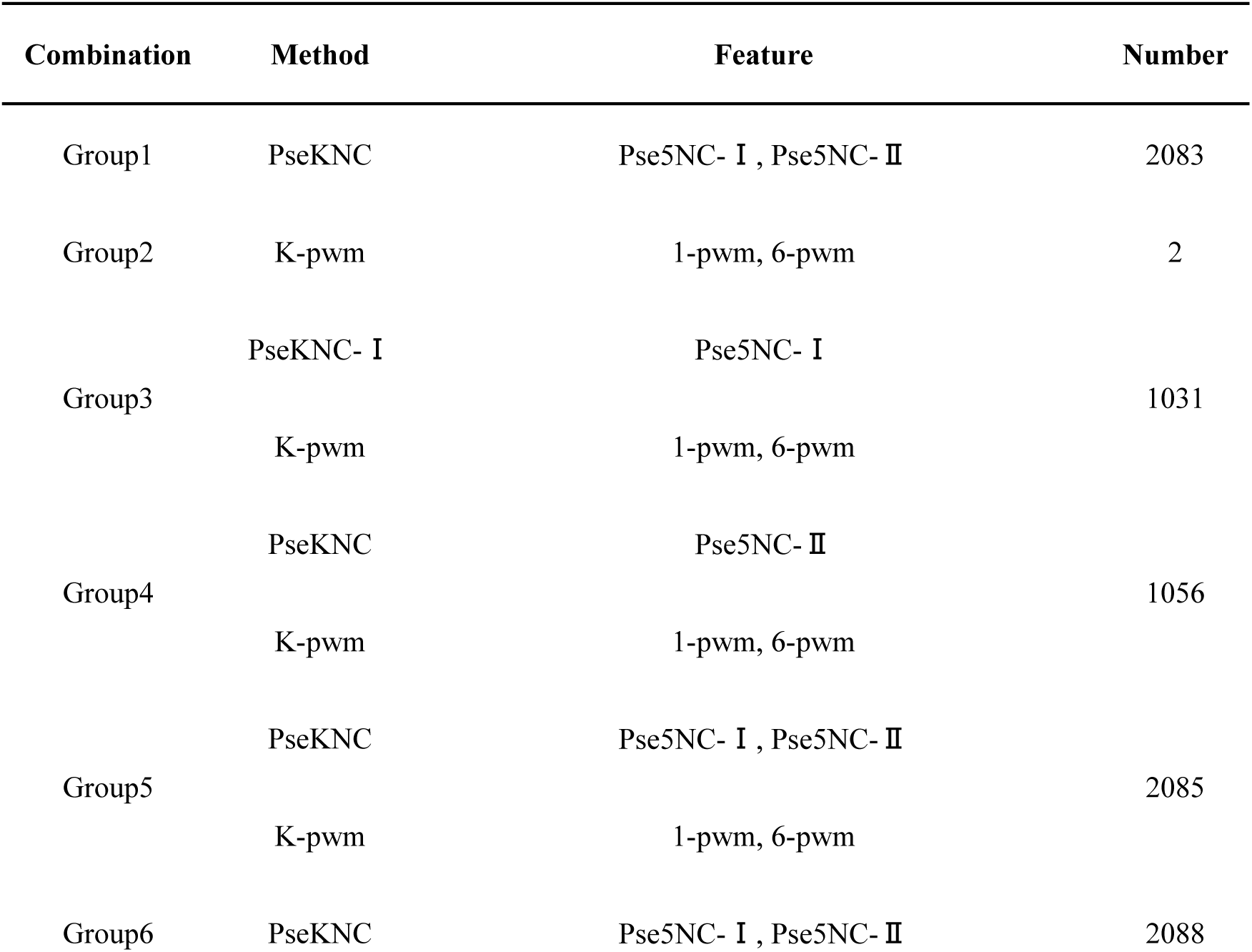
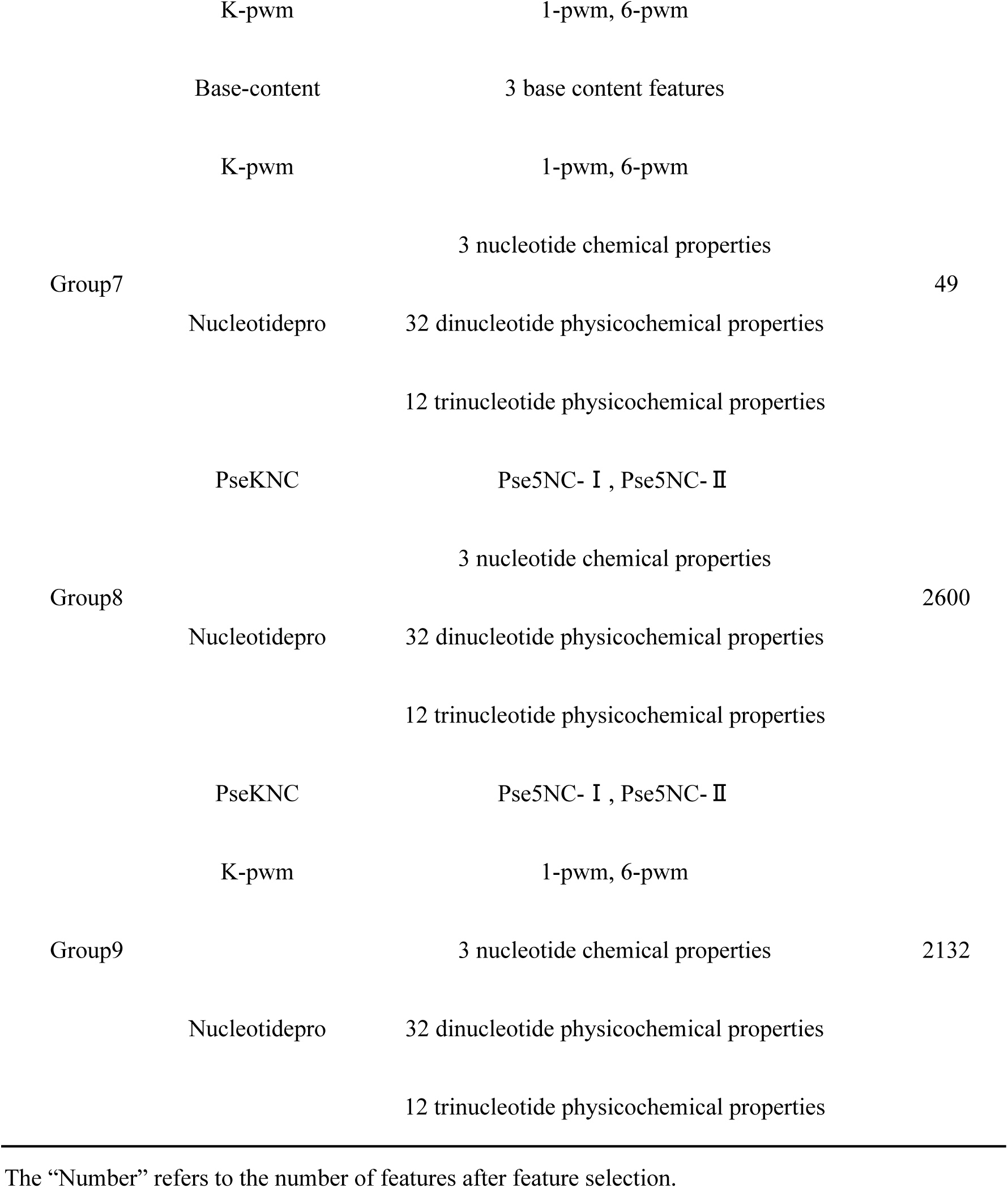
Combination of feature extraction methods.

**Fig 5.**
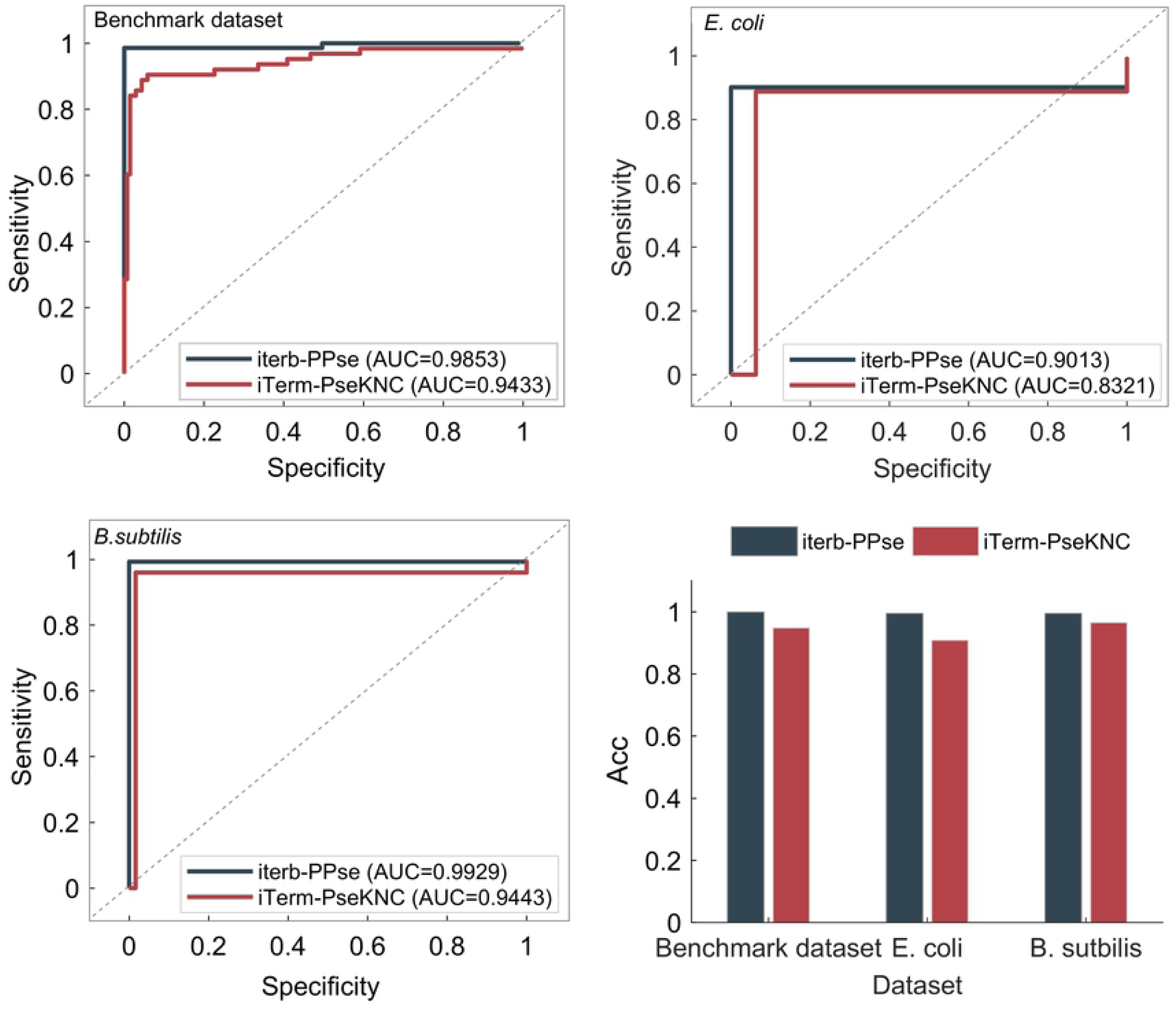
Prediction results using different feature extraction methods. All results are obtained after 100 times 5-fold CV. The ones marked red represent the best of each method.

**Fig 6.**
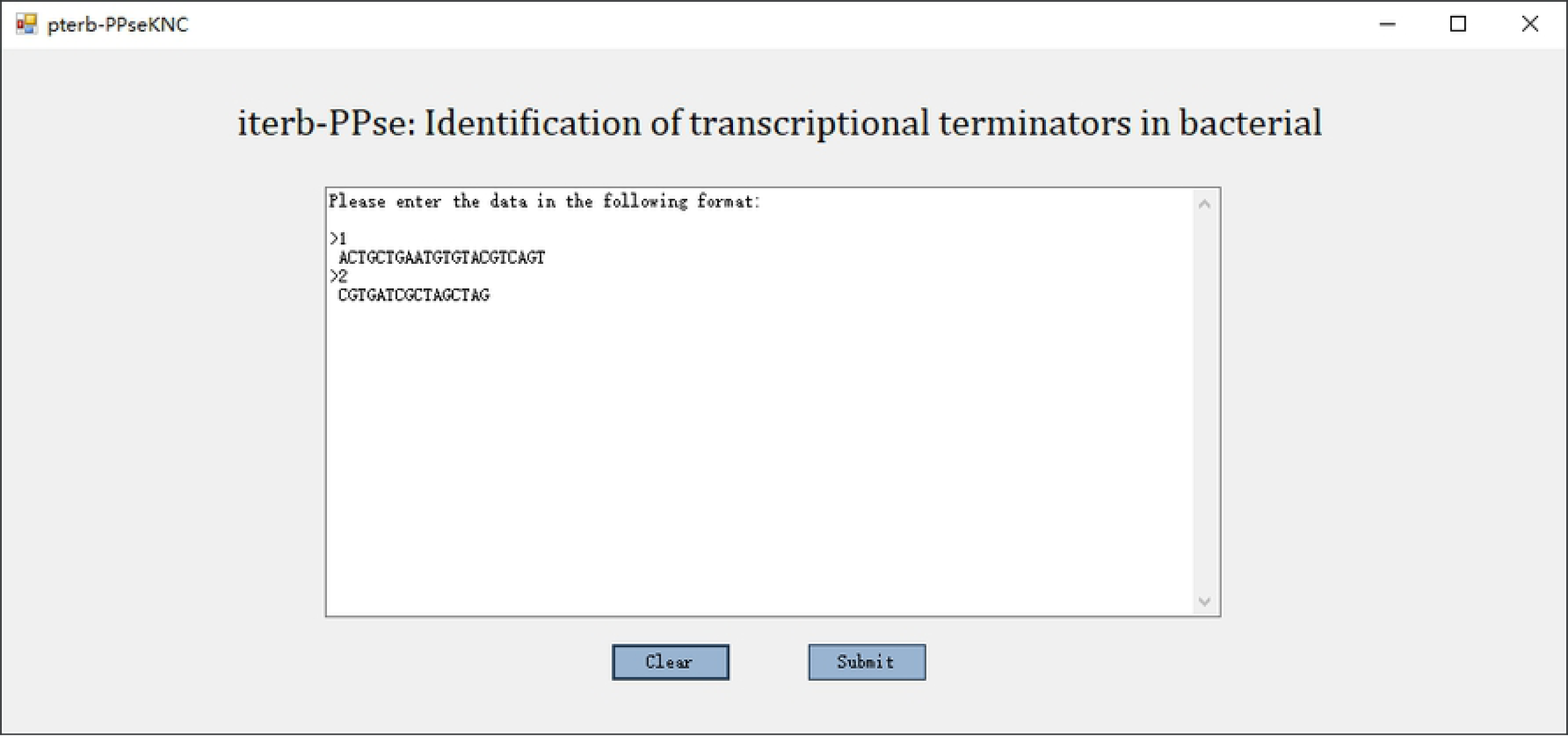
Classification results using different combined features. These results are obtained using XGBoost after 100 times 5-fold CV.

### 3.3 Comparison of different models

To compare different methods, the above experimental process was repeated using 16 different models. What can be clearly seen in Table 6 is that the classification performance of some ensemble models is better than that of a single model. For example, the accuracy of AdaBoost (SVM) and Bagging (SVM) are significantly higher than SVM. Decision tree, AdaBoost (Decision Tree) and XGBoost perform well, but XGBoost achieved the highest prediction accuracy in all models. Hence, it is reasonable and wise to choose XGBoost as the classifier.

**Table 6.**
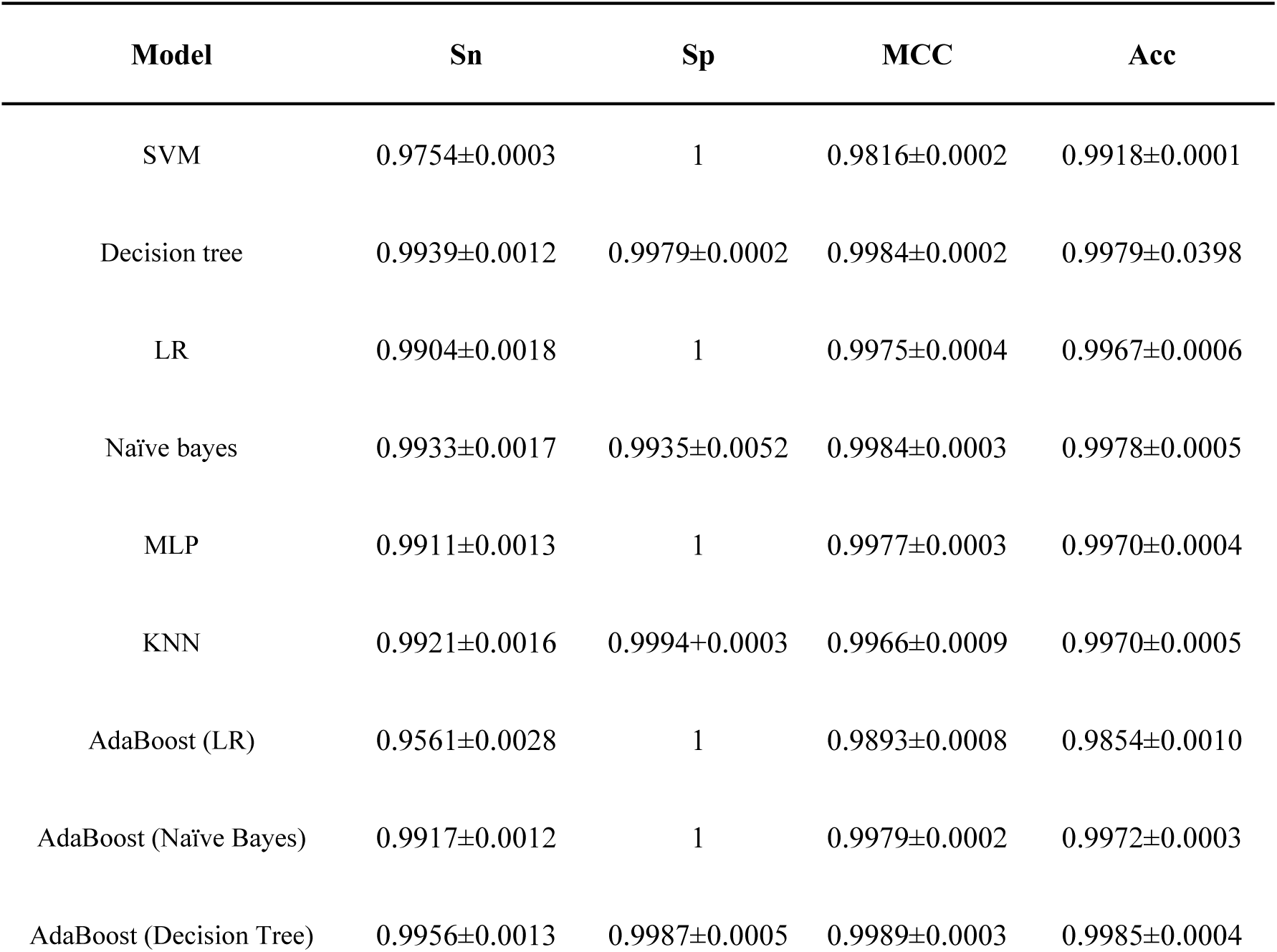
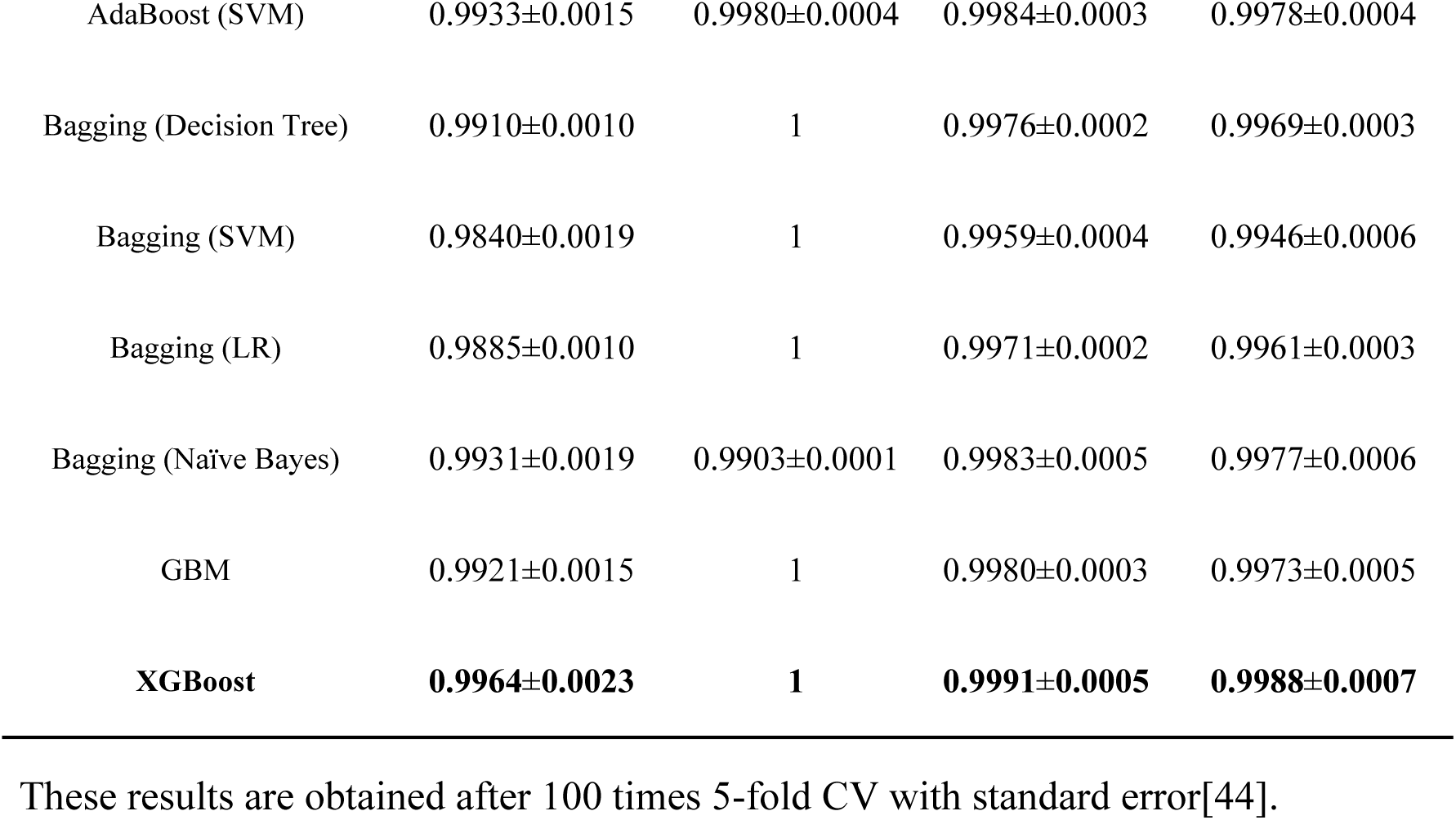
Display of all model classification results.

### 3.4 Comparison with existing state-of-the-art methods

To verify the advantage of our method “ iterb-PPse”, we made a comprehensive comparison with “ iTerm-PseKNC”[5], the current best tool for classifying two kinds of terminators, on the benchmark dataset and two independent sets we constructed using four evaluation parameters and ROC curves, as shown in Table 7 and Fig 7. The benchmark set we utilized is exactly the same with “iTerm-PseKNC”, so the comparison between the two methods is fair and objective.

**Table 7.** Comparison of “iTerm-PseKNC” and “iterb-PPse”.

| Dataset | Method | Sn | Sp | MCC | Acc |
| --- | --- | --- | --- | --- | --- |
| Benchmark dataset | iterb-PPse | **0.9964** | **1** | **0.9991** | **0.9988** |
|  | iTerm-PseKNC | 0.8545 | 0.9993 | 0.8846 | 0.9480 |
| *E. coli* | iterb-PPse | **0.9013** | **1** | **0.8898** | **0.9424** |
|  | iTerm-PseKNC | 0.8879 | 0.9371 | 0.8166 | 0.9084 |
| *B. subtilis* | iterb-PPse | **0.9929** | **1** | **0.9844** | **0.9945** |
|  | iTerm-PseKNC | 0.96 | 0.9836 | 0.9066 | 0.9653 |

The prediction results were obtained after 100 times 5-fold CV.

**Fig 7.**
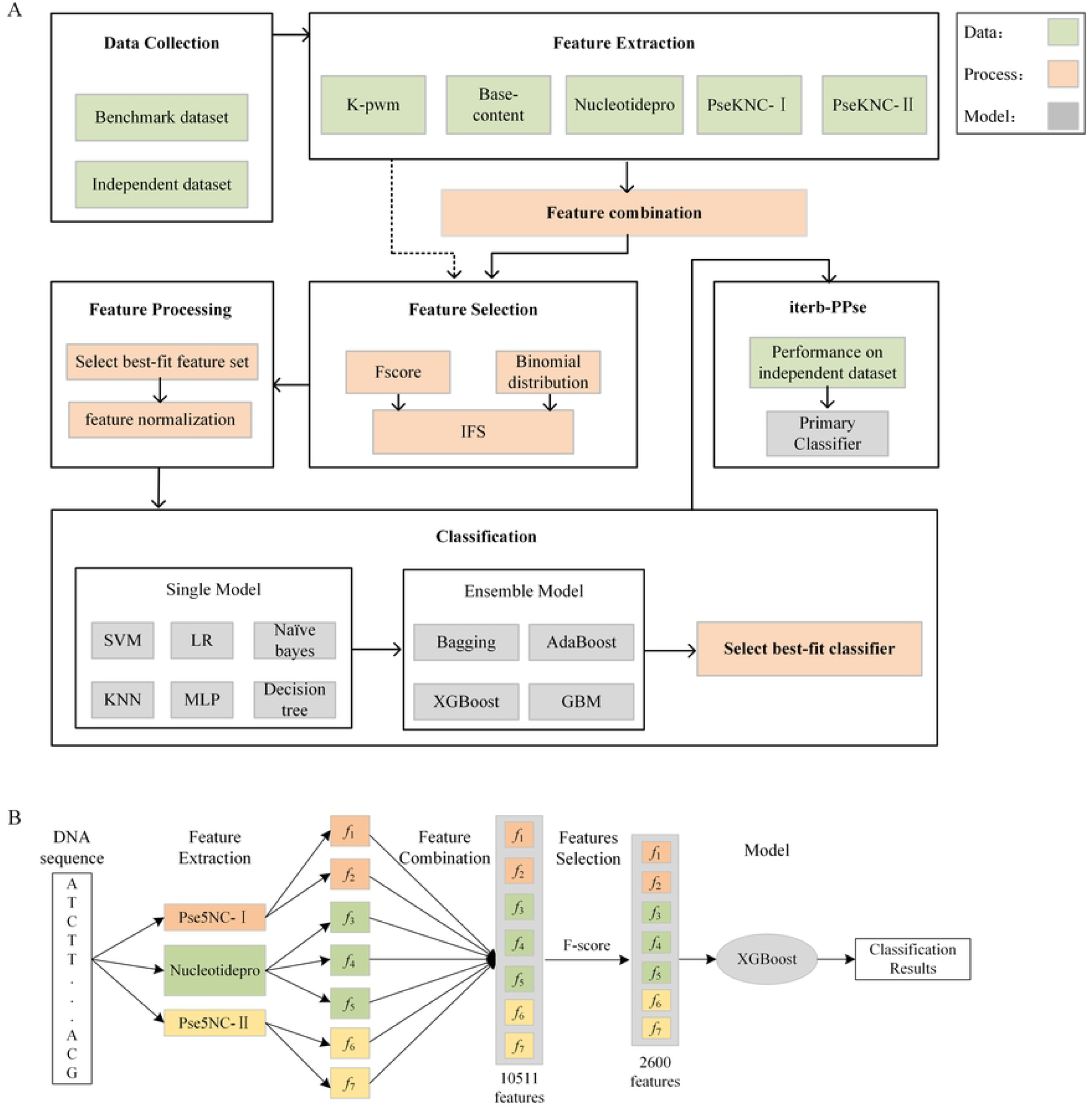
Comparison of “iTerm-PseKNC” and “iterb-PPse”. (A)-(C) ROC curves of two methods’ performance on the benchmark dataset and independent sets. (D) Prediction accuracy of two methods on different datasets.

As shown in Table 7 and Fig 7, the “iterb-PPse” is superior to the “iTerm-PseKNC” across the three datasets in Sn, Sp, MCC, Acc and AUC. Besides, the ROC curves in also show that the overall performance of our method is better. To be more precise, we improved the prediction accuracy (Acc) by 5.08%, 3.4%, 2.92% for the benchmark dataset and two independent datasets respectively.

### 3.5 Availability of software “iterb-PPse”

In addition to providing all codes of the prediction method, we developed a prediction software which could directly predict whether a DNA sequence is a terminator by simply installing it according to our software manual. The interface of the software is shown in the Figure 8.

**Figure 8.**
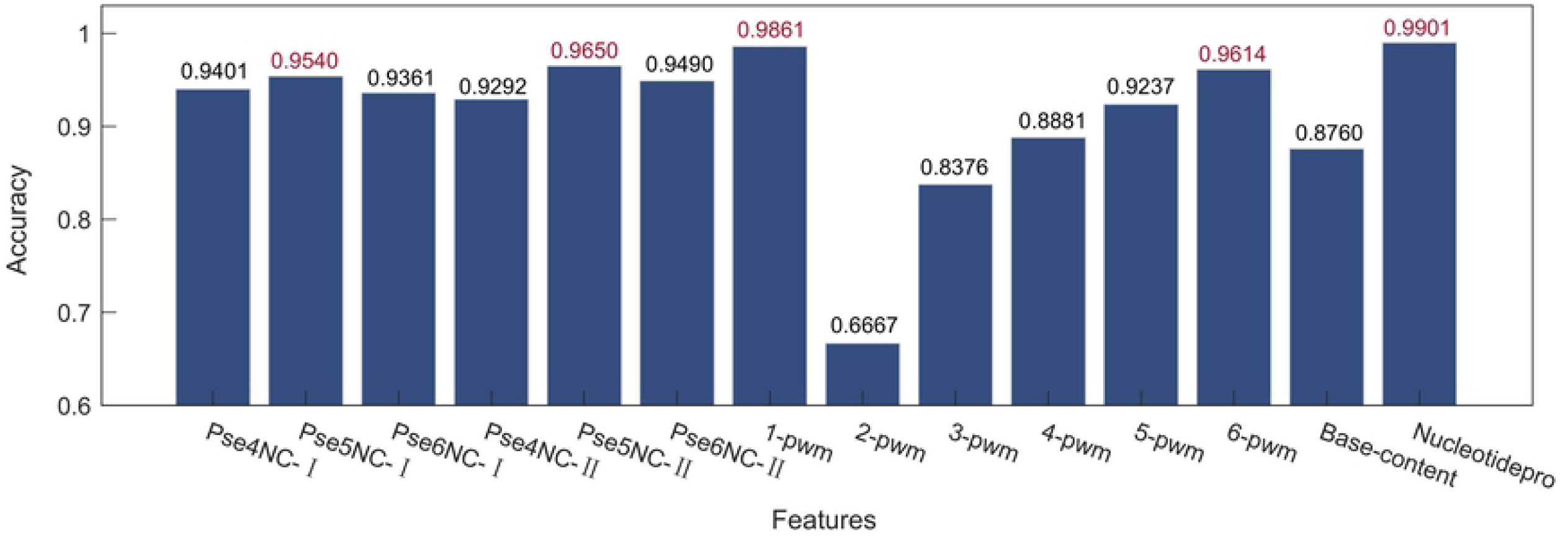
Main form of prediction tool. Just enter the sequence into the text box to get the prediction result.

## 4 Conclusions

In this work, we made miscellaneous comparisons of different feature extraction methods and models in many aspects. Eventually we proposed an accurate classification method “iterb-PPse” with 99.64%, 100%, 99.91% 99.88% in Sn, Sp, MCC, Acc respectively which is superior to the state-of-art prediction method and came to the following conclusions: (1) PseNC-`I`, PseNC-`II`, nucleotidepro are appropriate for formulating all samples. It proofs that nucleotide properties and the nucleotide components play a significant role in terminator classification and using the single GC content feature can not achieve the ideal classification effect. When using K-pwm feature extraction methods, we found that position-weight features of oligonucleotides and hexanucleotides are effective for predicting terminators (2) XGBoost works best on predicting terminators among all models based on the features we extracted. All the code and data used in our experiment are open source and are available at https://github.com/Sarahyouzi/myexperiment, hopefully could provide some assistance for related researches.

## Supporting information

**S1 Table. Dataset with 280 terminator sequences of *E. coli*.**

(CSV)

**S2 Table. Dataset with 560 non-terminator sequences of *E. coli*.**

(CSV)

**S3 Table. Dataset with 425 terminator sequences of *B. subtilis*.**

(CSV)

**S4 Table. Dataset with 147 terminator sequences of *E. coli*.**

(CSV)

**S5 Table. Dataset with 76 terminator sequences of *E. coli*.**

(CSV)

**S6 Table. Dataset with 159 non-terminator sequences of *E. coli*.**

(CSV)

**S7 Table. Dataset with 122 non-terminator sequences of *B. subtilis*.**

(CSV)

**S8 Table. Dinucleotide physicochemical properties.** This table contains 32 dinucleotide physicochemical properties we used and the corresponding standard values.

(CSV)

**S9 Table. Trinucleotide physicochemical properties.** This table contains 12 trinucleotide physicochemical properties we used and the corresponding standard values.

(CSV)

